# Prelimbic proBDNF facilitates memory destabilization by regulation of neuronal function in juveniles

**DOI:** 10.1101/2021.12.30.474526

**Authors:** Wei Sun, Xiao Chen, Yazi Mei, Yang Yang, Xiaoliang Li, Lei An

## Abstract

Fear regulation changes as a function of age and adolescence is a key developmental period for the continued maturation of fear neural circuitry. The involvement of prelimbic proBDNF in fear memory extinction and its mediated signaling were reported previously. Given the inherent high level of proBDNF during juvenile period, we tested whether prelimbic proBDNF regulated synaptic and neuronal functions allowing to influencing retrieval-dependent memory processing. By examining freezing behavior of auditory fear conditioned rats, we found high levels of prelimbic proBDNF in juvenile rats enhanced destabilization of the retrieval-dependent weak but not strong fear memory through activating p75^NTR^-GluN2B signaling. This modification was attributed to the increment in proportion of thin type spine and promotion in synaptic function, as evidence by facilitation of NMDA-mediated EPSCs and GluN2B-dependent synaptic depression. The strong prelimbic theta- and gamma-oscillation coupling predicted the suppressive effect of proBDNF on the recall of post-retrieval memory. Our results critically emphasize the importance of developmental proBDNF for modification of retrieval-dependent memory and provide a potential critical targeting to inhibit threaten memories associated with neurodevelopment disorders.

## 1. Introduction

Fear can be highly adaptive in promoting survival, yet it can also be detrimental when it persists long after a threat has passed (Kim-Cohen *et al*, 2003; Liberman *et al*, 2006; Moriceau and Sullivan, 2006). Flexibility of the fear response may be most advantageous during the early development period when animals are prone to explore novel, potentially threatening environments (Kim *et al*, 2011; McCallum *et al*, 2010; Pattwell *et al*, 2012). Actually, dynamic changes in neural circuitry during the sensitive period of neural development can induce memory to enter a destabilized state and attenuate maladaptive memories associated with emotional disorders (Pattwell *et al*, 2011; Pattwell *et al*, 2016). Retrieval of an existing fear memory can trigger either maintenance or inhibition of the stored memories depending on the degree of re-exposure to fear cues (Merlo *et al*, 2018). These opposing fear-related behaviors depend on distinct components of a medial prefrontal cortical circuit that receive projections from the amygdala and ventral hippocampus (Milad and Quirk, 2002; Sotres-Bayon *et al*, 2012). Further researches indicate that the developmental changes in the extinction and recall of cued fear memories involve protracted development of prelimbic region of medial prefrontal cortex (mPFC) (Da Silva *et al*, 2020; Kim *et al*, 2009). Importantly, the prelimbic subregion of mPFC remains immature after birth and develops, undergoing a process of programmed synaptic pruning followed by regulation of synaptic and neuronal function which can persist into adulthood and continue organization of maturation circuitry (Burgos-Robles *et al*, 2009; Sotres-Bayon *et al*, 2012).

Convergent adolescent rodent contextual fear findings further highlight the importance of the developing prelimbic cortex in mediating fear responses. For instance, during very early development in pre-weanling rodents, odor-shock conditioning can be modulated by maternal presence (Moriceau *et al*, 2006), and the mechanisms of cued extinction learning differ from adult-like extinction via alterations in N-methyl-D-aspartate (NMDA) receptors (NMDARs) requirement (Langton *et al*, 2007) and mPFC activity (Kim *et al*, 2009), suggestive of the perineuronal nets protect fear memories from degradation and erasure (Gogolla *et al*, 2009; Kim and Richardson, 2007). Potentiation of the prelimbic excitatory synapses after fear acquisition in postnatal day (PD) 23 mice, and its subsequent depotentiation upon extinction, suggest that the prelimbic region excitatory synapses dynamically regulate fear expression (Pattwell *et al*, 2012). Furthermore, the deep-layer prelimbic pyramidal neurons of juvenile animals can exert suppressive effects on fear-enhancing potentiation through the selective elimination of principally excitatory synapses (Pattwell *et al*, 2011; Pattwell *et al*, 2016). More convincingly, the mPFC shows enhanced functional connectivity during extinction of the reminded fear may play a role in enabling extinction learning training to more persistently modify the original threat-memory trace (Schiller *et al*, 2013). Therefore, understanding the underlying mechanism in memory destabilization upon reactivation may shed light on treatments for neurodevelopmental disorders, such as schizophrenia-spectrum and primary psychotic disorders, mood disorders and post-traumatic stress disorder (PTSD).

In stark contrast to adult animals, fear extinction of rats younger than 3 weeks appears to be permanent and has been suggested to reflect an unlearning process that leads to the erasure of previously conditioned fear memories (Kim and Richardson, 2008). Recent study has shown that the modification of an existing fear memory trace was associated with the retrieval-dependent alpha-amino-3-hydroxy-5-methyl-4-isoxazole propionate-type glutamate receptors (AMPARs) trafficking (Holehonnur *et* al, 2016). Fear circuitry during the retrieval of existing memories has been shown to be particularly plastic, which is a protein-synthesis-dependent process (Finnie and Nader, 2012; Lee, 2009; Tronson and Taylor, 2007). However, the neuronal mechanisms underlying the developmental regulation of retrieval-dependent memory are not fully known.

The precursor form of brain-derived neurotrophic factor (BDNF), proBDNF, which signals through its receptor p75^NTR^, has been shown to modulate structural plasticity, spine pruning and cell apoptosis during brain development. Abundantly proBDNF and its receptor are expressed in developing brain regions exhibiting a high degree of plasticity, such as the cerebellum, hippocampus and mPFC (Menshanov *et al*, 2015; Sun *et al*, 2021a; Yang *et al*, 2009c). During cortical development levels of proBDNF rise, the activation of proBDNF-p75^NTR^ signaling can negatively regulate neural remodeling by selectively facilitating NMDAR-dependent neurotransmission (Yang *et al*, 2014) and neuronal activity (Sun *et al*, 2019). Recently, we found that infusions of proBDNF in the infralimbic region facilitated induction of fear extinction, while infusions in the prelimbic region depressed fear expression (Sun *et al*, 2019; Sun *et al*, 2018a). This proBDNF-induced extinction is sufficient for extinguishing both new and older memories (Sun *et al*, 2018a). Indeed, prefrontal proBDNF is able to elevate neuronal correlate during the acquisition of conditioned extinction and facilitate dorsolateral striatum-dependent flexible behaviors (Sun *et al*, 2020b). Intriguingly, circadian proBDNF in the projection of the hippocampo-prefrontal cortex pathway possible engage in regulation of sleep homeostasis to extinguishing original memory (Sun *et al*, 2019). More recent research demonstrated that the peak expression of proBDNF during the 4^th^ postnatal week promoted the NMDAR-dependent short-lasting long-term depression (LTD), which was accompanied by inhibiting neuronal migration and axonal retraction (Sun *et al*, 2021a; Xu *et al*, 2011; Yang *et al*, 2009a). These retrograde effects of proBDNF on dendritic complexity and spine density are strongly linked to memory decay (Kasai *et al*, 2010).

Although previous reports provide important insights into pro-neurotrophic mechanisms for regulating fear extinction and weakening unwanted memory (Gibon *et al*, 2015; Yang *et al*, 2014; Yang *et al*, 2009c), it remains unclear whether and how prelimbic proBDNF regulate retrieval-dependent memory destabilization, especially when its high expression is observed during the early development stage. This study tested the hypothesis that the high expression of prelimbic proBDNF in the juvenile rodents influenced the modification of the weak fear memory traces. Furthermore, the experiments described here investigated the possible role of proBDNF in synaptic development and neuronal function via the p75^NTR^ and NMDAR signaling. These findings may help to understand the characterization of developmentally regulates synaptic and neuronal functions underlying inherently suppressing unwanted fearful memories.

## 2. Materials and Methods

### 2.1 Animals

All procedures were approved by the Care and Use of Animals Committee of Guizhou University of Traditional Chinese Medicine (SCXK-2013-0020). Juvenile (PD24-PD26) and adult (PD72-78) male Wistar rats (Beijing Vital River Laboratory Animal Technology Co., Ltd., China) were group-housed (two-four rat/cage) and reared under an environmentally controlled room (21±2°C; 45±5% humidity) and maintained in a 12-h light/dark cycle with food and water available *ad libitum*. To minimize possible circadian influences on learning and memory processing, all experiments were conducted during the light period (between 1400 and 1700). Handled extensively (10 min per day for at least two days) rats were averagely and randomly received one of infusion or co-infusion to excluded drug accumulation effects.

### 2.2 Behavioral manupulations

Using the same behavioral paradigm as in our previous studies, conditioned freezing behavior was tested in operant chambers (Coulbourn Instruments, Pennsylvania, USA) enclosed in ventilated sound-attenuating boxes (Med Associates Inc, Vermont, USA) with video system, which was supported by a video camera for scoring of animal’s behavior. Each chamber was illuminated by a dimly light and auditory stimuli were delivered from a programmable high-frequency speaker (ENV-224BM). Electrical footshock (2.0 s, 0.75 mA) was delivered through a grid floor via a constant current aversive stimulator (ENV-414S). The expression of fear behavior was automatically measured by the Med Associates software.

The procedure for four or eight tone-shock fear conditioning was referred a previously published model (Holehonnur *et al*, 2016; Wang *et al*, 2009). Rats were allowed to acclimate to the training apparatus for 150 s, followed by the presentation of a habituation tone (4 kHz, 30 s, 77 dB) with a footshock. Four tone-shock fear training used an ITI of 2 min and eight tone-shock fear training used a random but averaged 4 min ITI (range from 1 to 7 min). Rats were removed from the training chamber 30 s after the delivery of the last footshock and returned to their home cages. Fear memory test was performed one day after conditioning in a novel chamber with distinct environment (including tactile, visual, and olfactory cues). The test consisted of a 150 s acclimation period, followed by the presentation of four 30-s tone presentations with 2-min ITIs. Percentage freezing was measured during each tone presentation.

Five days after fear conditioning, rats were subjected to fear memory retrieval by exposing to a novel chamber with distinct environment (including tactile, visual, and olfactory cues). They were subjected to a 150 s acclimation period, were presented with one tone presentation for 30 s, and then were placed back to their home cage 30 s after the tone presentation. Percentage freezing was measured during the tone presentation. Post-retrieval short-term memory (PR-STM) and post-retrieval long-term memory (PR-STM) occurred 30 min and 2 days after the memory retrieval in a novel chamber with distinct cues, respectively. They were subjected to a 150 s acclimation period and four 30-s tone presentations with 2-min ITIs, and then were placed back to their home cage 30 s after the last tone presentation. Percentage freezing was measured during each tone presentation.

For fear extinction, rats were subjected to a 150 s acclimation period, followed by the presentation of eight 30-s tone presentations with 2-min ITIs five days after the fear conditioning. Percentage freezing was measured during each tone presentation. For the expression of fear, freezing behavior was reported as the freezing behavior during the first three tones. Fear reinstatement was conducted one day following the extinction training. During the reinstatement training, rats were subjected to two non-contingent footshock trials with a 1-min ITI. Percentage freezing was measured during each tone presentation.

### 2.3 Stereotaxic surgery and infusion

Rats were anesthetized with isoflurane and placed in a stereotaxic frame (SN-3, Narishige, Japan) for surgery. Stainless steel guide cannulae (22 gauge; Plastics One, Inc.) were bilaterally inserted above the prelimbic region of the mPFC (AP: +3.2 mm, ML: ±0.6 mm, DV: 3.2–3.6 mm). A sterile stainless steel stylet (30-gauge, 10 mm, Plastics One Inc.) was inserted into guide cannula to avoid obstruction.

During infusions, rats were gently restrained and infusions were achieved by inserting 30-gauge needles (10 mm, Small Parts Inc.) connected through PE-50 tube into a microsyringe pump (Harvard Apparatus), extended 1.0 mm beyond the end of the cannulae. Needles were inserted into bilateral cannulae and then cleavage-resistant proBDNF (2 ng/ml; Cat#B257 Alomone Labs), anti-proBDNF antibody (10 μg/μL; Cat#ANT-006, Alomone Labs), TAT-Pep5 (4 ng/μL; Cat#506181, EMD Millipore), K252a (25 μg/μL; Cat#82497; Sigma-Aldrich), 3-(2-carboxypiperazin-4-yl)propyl-1-phosphonic acid (CPP; 32 ng/μL; Cat#01773, Tocris Bioscience), NVP-AAM077 (0.8 ng/μL; Cat#P1999, Sigma-Aldrich), Ro25-6981 (2.0 ng/μL; Cat#1594, Tocris Bioscience), mature BDNF (1.5 μg/mL; Cat#B250; Alomone Labs) or artificial CSF (ACSF, Cat#3525, Tocris Bioscience) into prelimbic area (at a rate of 0.5 μL/min/side for 2 min) was infused immediately or one day following memory retrieval. The injection cannulae were left in place for 3-5 min before being slowly retracted. Dose and route of administration were chosen as described previously (An *et al*, 2018a; An and Sun, 2018b; Sun *et al*, 2018a). Rats were weighed and monitored at least three days to ensure recovery. One day before commencing experiment, a habituation session was performed three times for each rat without any infusion.

### 2.4 Tissue preparations and Western blot analysis

According to the previous principle (Sun *et al*, 2021c; Sun *et al*, 2021d), prelimbic cortex were dissected and homogenized in lysis buffer (pH 7.4) containing a cocktail of protein phosphatase and proteinase inhibitors (Sigma). The samples were centrifuged at 14,000 r.p.m. for 15 min at 4°C and the supernatant was collected. Protein was quantified and normalized to the bicinchoninic acid (BCA) assay. Protein amounts (20 μg) were resolved in 10-15 % SDS-PAGE gels followed by blotting onto polyvinylidene fluoride membranes (Pall, Pensacola). Samples were incubated with the primary antibody: mouse anti-proBDNF (1:500; Cat#sc-65514, Santa Cruz Biotechnology), rabbit anti-p75^NTR^(1:1,000; Cat#AB1554, Chemicon), rabbit antiGluN2A (1:1000; Cat#07632, Millipore), mouse anti-GluN2B antibody (1:1000; Cat#06600, Millipore), mouse anti-β-actin (1:20,000; Cat#A5316, Sigma) overnight at 4°C. After further incubation in horseradish-peroxidase (HRP)-conjugated secondary goat anti-mouse (1:2500; Cat#31430, Thermo Fisher Scientific) or anti-rabbit (1:2500; Cat#31460, Thermo Fisher Scientific) IgG for two hours at room temperature, immunoreactivity was detected by ECL Western Blotting Detection Kit (CWBIO, China). The relative values were calculated and normalized to the corresponding β–actin signal. The expression of the target proteins in juvenile group were presented as fold changes relative to the appropriate control values obtained from adult animals.

### 2.5 Spine density analysis

Rats were anesthetized by an intraperitoneal injection of sodium pentobarbital (80 mg/kg). The brains were removed, rinsed in phosphate-buffered saline (PBS), and stained using Golgi-Cox method, according to the manufacturer’s instructions (Rapid GolgiStain; FD Neurotechnologies). The brain was blocked and placed immediately into Golgi impregnation solution (a mixture of equal volumes of solution A and B provided in the kit) in the dark. The impregnation solution was replaced once after 24 hours and stored in the dark for two or three weeks at room temperature. Brain tissue was then transferred to a cryoprotectant solution (solution C) for one week at 4 °C. Tissues were frozen, embedded in OCT (ThermoFisher), and sectioned into 100 μm thick. Following air-drying in the dark, sections were rinsed with distilled water reacted in a developing solution, and dehydrated in 50%, 75%, 95%, and 100% ethanol for 5 min each. The slides were mounted under a coverslip with Permount and allowed to dry in the air. The region of interest was viewed and traced using a BX51 Olympus microscope (100×objective) attached to Olympus DP50 camera and reconstructed using Neurolucida 8.0 software (MBF Bioscience). All clearly evaluable areas of brain slices, which containing 50-150 μm of secondary dendrites from each imaged soma in layer V/VI were selected as previously described (Cerqueira *et al*, 2007; Moench and Wellman, 2017). Three pyramidal neurons per section and three sections per animal were analyzed. For spine categorization, the following criteria were used (Li *et al*, 2017; Sun *et al*, 2020b; Sun *et al*, 2021a): (1) mushroom: spine head diameter was ≥ 1.5 × spine neck diameter; (2) stubby: spine head and spine neck were roughly of same width, and spine length was not significantly longer than head diameter; (3) thin: spine head and spine neck were roughly of same width, and spine length was much longer than spine head width. Spine densities were calculated using NeuroExplorer software (MBF Bioscience) and expressed as the mean number of spines per micrometer dendrite.

### 2.6 Patch-clamp recording

As previously described (An and Sun, 2017; Li *et al*, 2018; Sun *et al*, 2021e), animals were decapitated and the brains were removed to an ice-cold, oxygenated (95% O_2_ and 5% CO_2_) high-sucrose solution that contained 200 mM Sucrose, 1.9 mM KCl, 1.2 mM NaH_2_PO_4_, 33.0 mM NaHCO_3_, 10.0 mM glucose, 4.0 mM MgCl_2_ and 0.7 mM CaCl_2_, pH 7.4 (with an osmolarity of 300–305 mOsm). Horizontal slices (300 μm thickness) were prepared with a vibratome (VT1000S, Leica, Germany). After a one hour recovery period, slices were recorded in a chamber mounted on a contrast-enhanced CCD camera (Hamamatsu) equipped with infrared gradient contrast. The slices were perfused with a continuous flow of ACSF (95% O_2_ and 5% CO_2_) that contained 120 mM NaCl, 3.5 mM KCl, 1.25 mM NaH_2_PO_4_, 26.0 mM NaHCO_3_, 10.0 mM glucose, 1.3 mM MgCl_2_ and 2.5 mM CaCl_2_, pH 7.4.

Whole-cell voltage-clamp recordings were performed in pyramidal neurons in layer V/VI of prelimbic region using pipettes with 3–7 M resistance after being filled with pipette solution containing (in mM) K-glu 130, MgCl_2_ 2, HEPES 10, EGTA 3, Na_2_-ATP 2, PH 7.3. The pipettes were pulled using a P-97 electrode puller (Sutter Instruments). All the cells were held at -70 mV, then slow and fast capacitance compensation was automatically performed. Access resistance was continuously monitored during the experiments. Neurons were considered only when the seal resistance was > 500 MΩ and the series resistance (< 30 MΩ) changed < 20% throughout the experiment. Positive pressure was applied to the recording pipette as it lowered into the medium and approached the cell membrane. Constant negative pressure was applied to form the seal (>1GΩ) when the recording pipette attached to the membrane. And then suck quickly to rupture the cell membrane and access whole cell configuration. All experiments were carried out at room temperature (22±1°C). Data of EPSC were recorded using an EPC-10 patch-clamp amplifier (HEKA Instruments). Signals were digitized at 10 kHz, low-pass filtered at 2.9 kHz, stored on a disk using Pulse software (HEKA, Germany). Synaptic spontaneous excitatory postsynaptic currents (sEPSCs) were isolated by bath application bicuculline (10 μM) to block the gamma-aminobutyric acid type A (GABA_A_) receptor mediated synaptic currents. Only one slice from each rat was used for final analyses. Spike curves were analyzed using Clampfit software (Molecular Devices). Decay time was calculated in Axograph X using a two-exponential simplex fit.

### 2.7 In vivo long-term depression

Thirty min after the memory retrieval or infusions, field excitatory postsynaptic potentials (fEPSPs) in at the hippocampal-prefrontal cortex pathway were recorded as published methods (Sun *et al*, 2020a; Takita *et al*, 2013; Thomases *et al*, 2014). Following craniotomies above the region of the mPFC and hippocampus, monopolar electrodes (insulated platinum iridium wire, AM Systems) and tungsten bipolar stimulating electrodes (FHC) were implanted into prelimbic region (AP: +3.2 mm, ML: ±0.6 mm, DV: 3.2–3.6 mm) and ventral hippocampus (AP: -6.0 mm, ML: ±5.2 mm, DV: 6.5-7.0 mm) regions, respectively. The cerebral hemisphere was chosen pseudorandomly for each animal. Neural signal was amplified (×100), filtered at 5-5000 Hz, digitized and collected at 20 kHz sample frequency (Scope software, PowerLab; AD Instruments). After optimal placement of the stimulating (100 ms pulse width, squared monophasic pulse) and recording electrodes, stimulus intensity was adjusted to about 50% of maximal fEPSP slope. Baseline fEPSPs were recorded every 60 s for at least 20 min. Long-term depression (LTD) was induced by a low-frequency stimulation (LFS; 300 pulses at 1.0 Hz). Initial data measurement was performed in Clampfit 9.0 (Molecular Devices). The fEPSPs slope was used to measure synaptic efficacy. The fEPSP slope at every time point was normalized to the mean fEPSP slope during the baseline period. Comparisons among the last 20 min of the 60-min decay were used to analyze.

### 2.8 LFP recording

Four or five days before behavioral training, microelectrode implantation was conducted using previously reported procedures (Sun *et al*, 2019; Sun *et al*, 2018b; Sun *et al*, 2021b). Rats were anesthetized with isoflurane during the surgery. An impedance-measured (200-600 kΩ) microelectrode, which was arrayed into a 4×8 matrix using 25-μm-diameter tungsten wires (California Fine Wires) in a 35-gauge silica tube (World Precision Instruments). A cannula was attached to a silica tube. The proximal open end of the cannula was parallel to electrode tips. The electrode array was chronically inserted into prelimbic region (AP: +3.2 mm, ML: ±0.6 mm, DV: 3.2-3.6 mm). Left or right hemisphere was implanted randomly but counterbalanced designed. The headmount was fastened to the cranium by dental acrylic with scull screws. A stainless steel wire was affixed to the skull to act as reference and ground electrodes.

Data were acquired on a Digital Cheetah system (Cheetah software, Neuralynx Inc.). Unit signals were recorded via a HS-36 unit gain headstage (Neuralynx Inc.) mounted on animal’s head by means of lightweight cabling that passed through a commutator (Neuralynx Inc.). LFPs were sampled at 32 kHz and filtered at 0.1–9,000 Hz from each electrode. Neural signals were transferred through a slip-ring commutator (Neuralynx) to the data acquisition system.

The LFPs during the memory retrieval were collected for further analysis, which was performed by using Neuroexplorer (Nex Technologies). The power of each frequency was calculated using Welch’s method (1024 frequencies between 1 and 200 Hz, smoothed with a Gaussian Kernel with bin width 3). The band power of theta and gamma were defined as the mean power in the frequency range of 6-12 Hz and 50-80 Hz, respectively.

To quantify the amplitude modulation by phase, Tort’s modulation index (MI) was calculated using a normalized entropy measure previously described (Sun *et al*, 2018b; Tort *et al*, 2008; Voloh *et al*, 2015). Briefly, the preprocessed voltage trace from a single trial, *x*_*raw*_*(t)* (raw signal), was filtered into two frequency bands of interest *f*_*P*_ and *f*_*A*_, generating *x*_*fP*_*(t)* and *x*_*fA*_*(t)*, respectively. The standard Hilbert transform was applied to extract the time series of the phase *φ*_*fP*_*(t)* and amplitude envelope *A*_*fA*_*(t)* from *x*_*fP*_*(t)* and *x*_*fA*_*(t)*, respectively. The composite time series [*φ*_*fP*_*(t), A*_*fA*_*(t)*] were then constructed, which calculated the amplitude of the *f*_*A*_ phase onto each phase of the *f*_*P*_ rhythm. The phase *φ*_*fP*_*(t)* were binned into interval of phase (*n*=18; 20 intervals). The mean amplitude 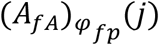 of *A*_*fA*_*(j)* was computed over each phase bin *j*. The entropy measure *H* was calculated as follows: 

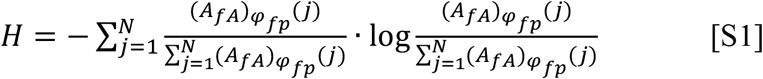

Where *N*=18, which was equal to the number of bins.

The null hypothesis of the test is that the expected amplitude distribution is uniform. Thus, the MI is the normalized H by the uniform distribution (log*(N)*),

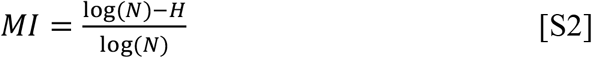

The range of MI value was larger than zero. A value above zero indicated the phase-to-amplitude modulation. The stronger phase-to-amplitude modulation, the larger MI value would be.

### 2.9 Statistical analysis

Data are expressed as mean ± S.E.M.. All analyses were performed with Neuroexplorer and SPSS 17.0 software. One-Sample Student t-tests were used to compare data from western-blot, spine density and fear expression. Other data were analyzed with one-way or two-way repeated analysis of variance (ANOVA). Post hoc analyses were performed with Tukey’s test where appropriate. *P*<0.05 level of confidence were used in the analyses. The number of animals in each group and other details can be found in figure legends and results.

## 3. Results

### 3.1 The increase proBDNF during the juvenile period enhances retrieval-dependent fear memory destabilization

The expression of proBDNF and its receptor p75^NTR^ in the prelimbic region of mPFC were detected in un-manipulated animals. Both proBDNF (Figure 1A; T-test, *t*_10_=2.54, *P*=0.030) and p75^NTR^ (Figure 1B; T-test, *t*_10_=2.32, *P*=0.043) levels of juvenile rats are significantly higher than adults. To investigate whether different intensity of auditory fear conditioning affect memory strength, rats were fear conditioned with either four tones or eight tones. The eight-tone groups of both juvenile and adult rats exhibited significantly higher tone-induced freezing than the four-tone groups (Figure 1C; two-way ANOVA, effect of tone: *F*_(1, 18)_=5.39, *P*=0.032) without age differences (effect of age: *F*_(3, 18)_=0.13, *P*=0.941), indicating that the eight-tone training resulted in stronger memories. To determine whether the increase proBDNF during the juvenile period was involved in fear memory destabilization, rats were bilaterally infused with proBDNF (Adult+Pro, Juvenile+Pro), anti-proBDNF antibody (Adult+Anti, Juvenile+Anti), anisomycin (Juvenile+Anis) or vehicle (Adult, Juvenile) immediately following memory retrieval. For rats trained with four tones, repeated-measures ANOVA indicated no statistical difference in freezing levels among groups during memory retrieval (Figure 1D). However, there was a significant decrease in freezing in the juvenile group compared with adult group during PR-LTM test (repeated-measures ANOVA, interaction effect of between treatment and test: *F*_(5, 31)_=4.21, *P*=0.005; post-hoc, Juvenile or Adult, *P*<0.05), which was conducted two days after memory retrieval. For the juvenile groups, infusion proBDNF or anisomycin depressed freezing behavior (post-hoc, Juvenile vs. Juvenile+Anis or Juvenile+Pro, both *P*<0.05), while infusion anti-proBDNF antibody enhanced the freezing levels (post-hoc, Juvenile or Juvenile+Anti, *P*<0.05). However, proBDNF-infused adult rats did not display less freezing (post-hoc, Adult vs. Adult+Pro, *P*>0.05). Therefore, endogenous proBDNF of juvenile rats facilitated retrieval-dependent destabilization, while exogenous proBDNF could regulate the memory destabilization of juvenile but not adult animals. For rats trained with eight tones, repeated-measures ANOVA revealed no statistical difference in freezing levels among groups during memory retrieval or PR-LTM test (Figure 1E; repeated-measures ANOVA, interaction effect of between treatment and test: *F*_(5, 24)_=0.22, *P*=0.951). Consistent with previous reports (Holehonnur *et al*, 2016; Wang *et* al, 2009), these findings imply that strong memories formed via eight-tone training do not undergo retrieval-dependent memory destabilization. To investigate whether the memory destabilization was resulted from changes in PR-STM, post-retrieval tests were conducted in rats trained with four tones 30 min following the memory retrieval. There was no interaction effect of between treatment and test on the freezing level (Figure 1F; repeated-measures ANOVA, interaction effect of between treatment and test: *F*_(5, 20)_=0.18, *P*=0.967). Meanwhile, proBDNF, its anti-body or anisomycin effects on the freezing behavior during the PR-LTM tests were not observed when juvenile rats trained four tones were infused one day after memory retrieval (Figure 1G; repeated-measures ANOVA, interaction effect of between treatment and test: *F*_(5, 20)_=2.93, *P*=0.038; post-hoc, Juvenile vs. Juvenile+Pro or Juvenlie+Anti or Juvenile+Anis, all *P*>0.05), suggesting there was no remote effect of proBDNF or anisomycin on post-retrieval memory during the juvenile period. Similarly, no significant difference was found between Adult and Adult+Pro group during memory retrieval (post-hoc, Adult vs. Adult+Pro, *P*>0.05) or PR-LTM test (post-hoc, Adult vs. Adult+Pro, *P*>0.05). Together, these results indicate the high expression of endogenic proBDNF in the juvenile rat prelimbic cortex plays a central role in promoting retrieval-dependent memory destabilization of a weak fear memory trace.

**Figure 1.**
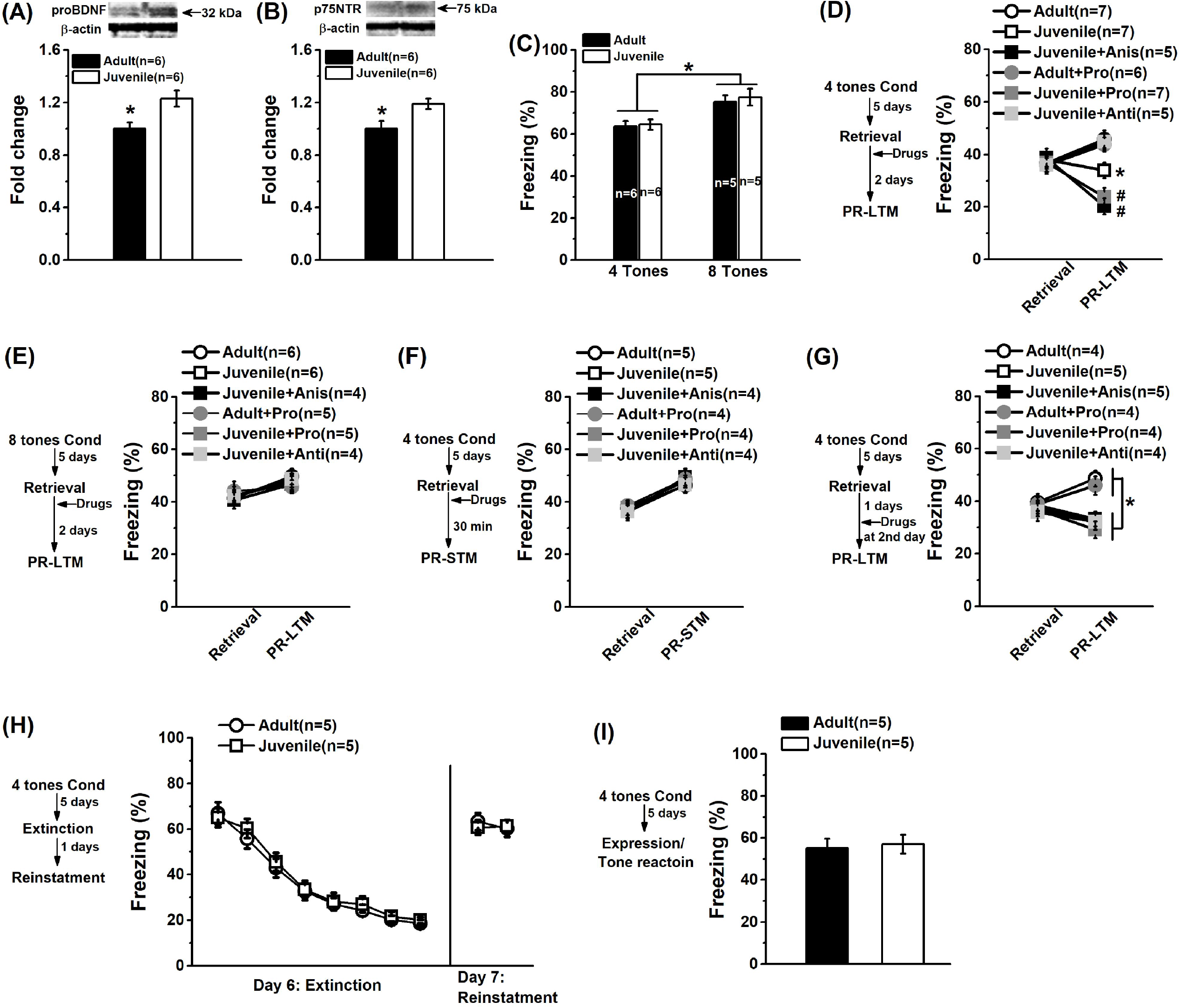
The higher prelimbic proBDNF expression during the juvenile period facilitates retrieval-dependent memory destabilization. Quantification of the proBDNF **(A)** and its receptor p75^NTR^ **(B)** levels in prelimbic cortex of juvenile and adult rats. Representative immunoblots the expression of proBDNF and p75^NTR^ (Top). A significant increase in the proBDNF levels was detected in juvenile group, as well the p75^NTR^ levels (Bottom). (\**P*<0.05). **(C)** Both juvenile and adult rats conditioned with eight tones froze significantly more than rats trained with four tones during the LTM test. (\**P*<0.05, four-tone group vs. eight-tone group). **(D)** Schematic describing the behavioral timeline for the retrieval-dependent memory destabilization experiment using rats conditioned with four tones (left). Rats were auditory fear conditioned with four tones. Retrieval of the fear memory was induced by exposing the rats to a single tone presentation five days following the fear condition learning, and then the rats were immediately infused with proBDNF, its antibody, anisomycin or vehicle into the prelimbic cortex. Two days later, the rats were exposed to tones in a novel context during the PR-LTM test and freezing was measured. No significant difference in the average percentage freezing across four tones during the memory retrieval but the percentage freezing during the PR-LTM test was significant lower in juvenile group than adult group (Right). Infusions of anti-proBDNF antibody obviously enhanced the freeze behavior of juvenile but not adult rats. Microinjections of anisomycin and proBDNF into the prelimbic region could enhance the freezing levels of juvenile rats. (\**P*<0.05, Juvenile group vs. other groups; #*P*<0.05, Juvenile+Pro or Juvenile+Anis group vs. other groups). **(E)** Schematic describing the behavioral timeline for the retrieval-dependent memory destabilization experiment using rats conditioned with eight tones (left). No significant difference in either the average percentage freezing across eight tones during the memory retrieval or the average percentage freezing during the PR-LTM test, which was conducted two days later (Right). **(F)** Schematic describing the experimental timeline for a reconsolidation experiment using rats conditioned with four tones and a PR-LTM test conducted 30 min later (left). No significant difference in either the average percentage freezing across four tones during the memory retrieval or the average percentage freezing during the PR-LTM test (Right). **(G)** Schematic describing the experimental timeline for a reconsolidation experiment using rats conditioned with four tones and drugs were infused one day after the memory retrieval (left). No significant difference in either the average percentage freezing across four tones during the memory retrieval but infusions of anisomycin and proBDNF did not enhance the freeze behavior during the PR-LTM test, which was performed one day after the microinfusions (Right). **(H)** Schematic describing the experimental timeline for fear extinction and reinstatement using rats conditioned with four tones (left). Rats were fear conditioned with four tones, and five days later, fear extinction learning was performed. The extinguished fear was reinstated one day following memory extinction. Juvenile group was indistinguishable from adult group in the acquisition of fear extinction training (Right). **(I)** Schematic describing the experimental timeline for fear expression (left). For tone reactions and the expression of fear behavior, the mean freezing levels during the first three tones of the extinction training were calculated. There was no significant difference in the percentage freezing (Right). The number of rats in each group used was indicated in each column/figure.

Rats trained with four tones were subjected to fear extinction training five days after fear conditioning. There was no significant main effect of age on acquisition of fear memory extinction (Figure 1H; repeated-measures ANOVA, effect of age: *F*_(1, 8)_=0.017, *P*=0.898). The inhibited freezing levels of both juvenile and adult groups could be completely regained when a reinstatement training was conducted one day following fear extinction (repeated-measures ANOVA, effect of age: *F*_(1, 8)_=0.021, *P*=0.889). Together, these findings imply that the high level of proBDNF expression during the juvenile period did not inhibit fear extinction or expression, or degrade the original fear memory. Meanwhile, there was no statistical difference in the mean fear expression of the first three tones during extinction training (Figure 1I; T-test, *t*_8_=0.19, *P*=0.858). We also ruled out the possible difference in locomotion, anxiety or motivation between juvenile and adult as rats had a similar travel distance (Supplementary Figure S1B) and the proportion time in the center field (Supplementary Figure S1C) during the open field test, and a similar motivation behavior in the lever-press task (Supplementary Figure S1D). Therefore, the performance of juvenile rats during the PR-LTM test is not attributed to the alteration of locomotion, anxiety or motivation.

### 3.2 The proBDNF/p75^NTR^ signaling mediated by the activation of GluN2B is involved in retrieval-dependent memory destabilization

To further confirm if the proBDNF-mediated memory destabilization involved in p75^NTR^ or TrkB pathway, p75^NTR^ (TAT-Pep5) or TrkB (K252a) inhibitor was bilaterally infused immediately following memory retrieval. Blocking prelimbic p75^NTR^ by the TAT-Pep5 inhibitor significantly enhanced the freezing level during the PR-LTM test (Figure 2A; repeated-measures ANOVA, interaction effect of between group and day: *F*_(5, 23)_=3.34, *P*=0.021; post-hoc, Juvenile(Con+Con) vs. Juvenile(Pep5+Con), *P*<0.05). Meanwhile, the effects of exogenous proBDNF could be prevented by administration of TAT-Pep5 (post-hoc, Juvenile(Con+Pro) vs. Juvenile(Pep5+Pro), *P*<0.05), but not K252a (post-hoc, Juvenile(Con+Pro) vs. Juvenile(K252a+Pro), *P*<0.05), indicating that retrieval-dependent memory destabilization was associated with the proBDNF/p75^NTR^–mediated pathway. To detect whether neutralizing proBDNF with its antibody would potentially interfere with the expression of endogenous protein and its proteolysis to mBDNF, rats were infused with mBDNF following memory retrieval. The freezing level of mBDNF-infused rats was similar to vehicle group during the PR-LTM test (post-hoc, Juvenile(Con+Con) vs. Juvenile(Con+mBDNF), *P*>0.05). These observations following the infusion reflected merely the proBDNF effect rather than the mBDNF effect. The findings clearly indicate that proBDNF-p75^NTR^ signaling influences the modification of an existing memory trace.

**Figure 2.**
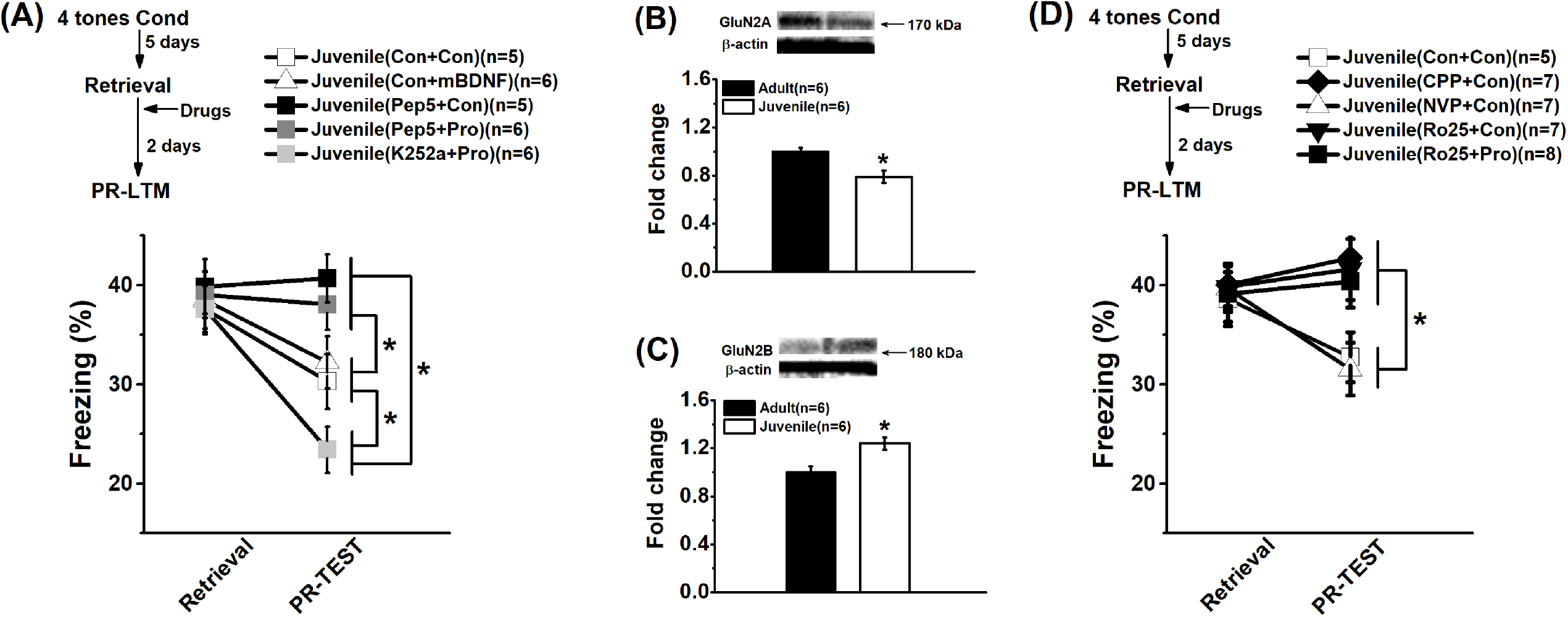
Up-regulation of proBDNF-p75NTR signaling mediated by NMDA-GluN2B contributes to enhance the modulation of existing fear memory traces in juvenile rats. **(A)** Schematic describing the behavioral timeline for the retrieval-dependent memory destabilization experiment using rats conditioned with four tones (Top). Immediately following the memory retrieval, the rats infused with TAT-Pep5, K252a or vehicle into the prelimbic cortex 15 min prior to the mBDNF, proBDNF or vehicle infusion. Two days later, PR-LTM was assessed by exposed the rats to the novel context. Similar, no significant difference in the percentage freezing during the memory retrieval but the percentage freezing level during the PR-LTM test was significant lower in juvenile group than adult group (Bottom). No obvious effect of mBDNF on freeze behavior was found. Infusions of p75^NTR^ blocker TAT-Pep5 could significantly enhance the percentage of freeze behavior. Meanwhile, infusions of TAT-Pep5, but not K252a, markedly blocked the effects of proBDNF treatment. (\**P*<0.05). Representative immunoblots the expression of the expression of the NMDA-GluN2A **(B)** and NMDA-GluN2B **(C)** receptors in prelimbic cortex (Top). The level of GluN2A subunit was significantly higher but the level of GluN2B subunit was significantly lower compared adult group with juvenile group. (\**P*<0.05). **(C)** Schematic describing the behavioral timeline for the retrieval-dependent memory destabilization experiment using rats conditioned with four tones (Top). Immediately following the memory retrieval, the rats infused with CPP, NVP-AAM077, Ro25-6981 or vehicle into the prelimbic region 15 min prior to the proBDNF or vehicle infusion. Two days later, the PR-LTM was tested as above. Both the broad NMDA receptor inhibitor CPP and The NMDA-GluN2B subunit antagonist Ro25-6981 enhanced the percentage freeze level during the PR-LTM test. However, the NVP-infused rats could maintain the lower percentage freeze level. Furthermore, blockage of the NMDA-GluN2B subunit could completely block the effects of proBDNF on freeze behavior. The number of rats in each group used was indicated in each column/figure.

Growing evidences suggest that NMDAR activity at the very moment of memory retrieval play an important role in updating reconsolidation and inducing memory destabilization (Ben Mamou *et al*, 2006; Holehonnur *et al*, 2016; Milton *et al*, 2008). Meanwhile, previous studies indicated that BDNF-induced fear extinction and memory consolidation were linked with the activation of NMDA receptors (Ma *et al*, 2021; Sotres-Bayon *et al*, 2009; Sun *et al*, 2019; Sun *et al*, 2018a). Furthermore, depending on the age of the animals, the dynamic changes in the expression of GluN1, GluN2A, and GluN2B subunit mRNAs can lead to different mixtures of NMDA receptors during development of rat cortex (Liu *et al*, 2004; Ohno *et al*, 2010; Sheng *et al*, 1994). To determine whether the proBDNF-mediated memory destabilization was regulated by NMDA receptors, we first examined the levels of NMDA-GluN2A and NMDA-GluN2B receptors in prelimbic cortex of both adult and juvenile rats. Then juvenile rats were subjected to four-tone fear conditioning as above, followed by infusion of NMDA (CPP), NMDA-GluN2A (NVP-AAM077) or NMDA-GluN2B (Ro25-6981) receptor antagonist into the prelimbic region immediately after memory retrieval. The NMDA-GluN2A level of juvenile group was significant lower than adult group (Figure 2B; T-test, *t*_10_=2.86, *P*=0.017) while a higher level of NMDA-GluN2B in the juvenile group was found (Figure 2C; T-test, *t*_10_=2.69, *P*=0.019). Both CPP and Ro25 enhanced the fear behavior during PR-LTM test (Figure 2D; repeated-measures ANOVA, interaction effect of between group and test: *F*_(5, 29)_=2.87, *P*=0.032; post-hoc, Juvenile(Con+Con) vs. Juvenile(CPP+Con) or Juvenile(Ro25+Con), both *P*<0.05) but NVP-treated rats exhibited a similar freezing level to vehicle rats (post-hoc, Juvenile(Con+Con) vs. Juvenile(NVP+Con), *P*>0.05). Furthermore, the proBDNF effects on memory destabilization could be effectively blocked by GluN2B antagonist Ro25 (post-hoc, Juvenile(Con+Con) vs. Juvenile(Ro25+proBDNF), *P*<0.05). These results demonstrate the involvement of proBDNF-NR2B pathway in proBDNF-mediated the induction of reconsolidation updating.

### 3.3 The changes in synaptic and neuronal function promote the modification of the existing fear memory trace

Previous evidence supports that proBDNF signaling plays a critical role in synapse formation during early brain development (Je *et al*, 2012; Langlois *et al*, 2013; Orefice *et al*, 2016; Sun *et al*, 2021a). Furthermore, dendritic spine constitutes the main locus of excitatory synaptic interaction among central neurons. The great variety of shapes and abundance has focused considerable attention on the dendritic spine as the key site for the potential encoding of activity-dependent, neuronal plasticity (Engert and Bonhoeffer, 1999; Weber *et al*, 2016). Given the relation between proBDNF and synaptogenesis during early brain development, it is important to evaluate the proportion of dendritic spine, neuronal and synaptic function during the early postnatal period, which may help understand the mechanism of retrieval-dependent memory destabilization.

Six hours after the memory retrieval, prelimbic cortex was collected and analyzed alterations in dendritic morphology. For the total density of dendritic spines, no significant difference was observed between juvenile and juvenile+anti rats (Figure 3A(bottom-left); T-test, *t*_10_=0.27, *P*=0.403). However, the percentage of mushroom spine types was significant lower in the juvenile group in comparison to the anti-proBDNF group (Figure 3A(bottom-right); T-test, *t*_10_=2.29, *P*=0.047) while the proportion of thin spines of juvenile rats was significant higher than juvenile+anti group (Figure 3A(bottom-right); T-test, *t*_10_=2.33, *P*=0.042). No statistical difference was found in stubby spines between two groups (Figure 3A(bottom-right); T-test, *t*_10_=0.13, *P*=0.901). There was no difference in the mean frequency of NMDA currents (Figure 3B-(bottom-left); one-way ANOVA, *F*_(2, 15)_=1.26, *P*=0.312). However, juvenile rats exhibited significant higher NMDA-EPSC amplitude than adults (Figure 3B-(bottom-middle); one-way ANOVA, *F*_(2, 15)_=5.72, *P*=0.014; post-hoc, Juvenile vs. Adult, *P*<0.05) while proBDNF-inactivating antibody could effectively depressed the amplitude of juvenile group (post-hoc, Juvenile vs. Juvenile+Anti, *P*<0.05). The decay time of NMDA receptor currents was significantly enhanced in juvenile rats compared with adult rats (Figure 3B-(bottom-right); one-way ANOVA, *F*_(2, 15)_=4.95, *P*=0.022; post-hoc, Juvenile vs. Adult, *P*<0.05). Blockage of proBDNF significantly shortened the decay time of juvenile group (post-hoc, Juvenile vs. Juvenile+Anti, *P*<0.05), observing no difference between Adult and Juvenile+Anti groups (post-hoc, Adult vs. Juvenile+Anti, *P*>0.05).

**Figure 3.**
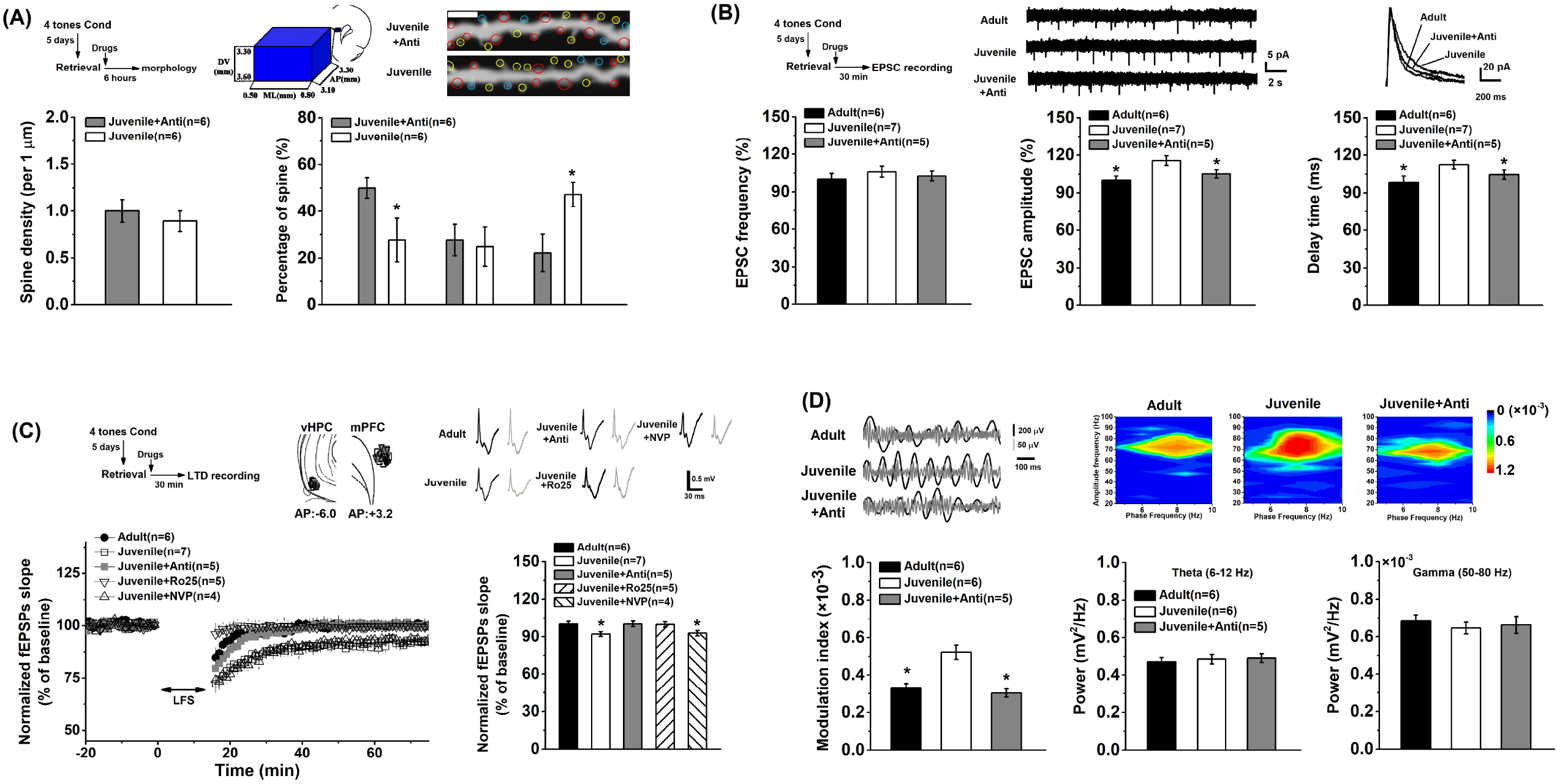
The increase proBDNF alters synaptic currents, promotes LFS-induced synaptic depression and strengthens the theta phase-gamma amplitude coupling during the PR-LTM test. **(A)** Schematic describing the timeline for morphological analysis (Top-left). Illustration of the region of interest in prelimbic images (Top-middle), and dendritic segment analysis for spine quantification (Top-right). Red circles indicated the mushroom type spine, yellow circles indicated thin type spine and blue circles indicated stubby type spine. Sample images were projected at minimal intensity and inverted, background was then subtracted, followed by brightness/contrast adjustment. Scale bars, 5 μm. Quantification of spine density (Bottom-left) and the proportion of spine (Bottom-right). No statistical difference in spine density was found between juvenile and adult groups. However, a significant higher proportion of thin type spine but a lower mushroom type spine was observed in juveniles compared with adults. **(B)** Schematic describing the timeline for EPSCs recordings (Top-left). Representative continuous traces (Top-middle) and average waveform (Top-right) of the pharmacologically isolated NMDA EPSCs in the prelimbic neurons of adult, juvenile and juvenile+anti groups. No change in the amplitude of EPSCs (Bottom-left) was found but the frequency (Bottom-middle) and decay time (Bottom-right) were significantly increased in juvenile group. The enhanced frequency and decay time of NMDA currents in juvenile group were inhibited after infusions of anti-proBDNF antibody. (\**P*<0.05, Adult or Juvenile+Anti group vs. Juvenile group). **(C)** Schematic describing the timeline for the LFS-induced LTD recordings (Top-left). Schematic representative placements of the stimulation electrodes (vHPC; ventral hippocampus) and recording electrodes (mPFC) (Top-middle). Numbers in figures were referenced to bregma. The black traces indicated the baseline traces and the dark gray ones indicated the traces of the fEPSPs, which were taken at 41 min after LFS (Top-right). Characteristic time courses of fEPSP slope (Bottom-left). Bidirectional arrow represented application of LFS (300 pulses at 1.0 Hz). Time coursing changes in fEPSPs slope. Magnitude of LTD was determined as responses in the last 20 min (between 41 and 60 min following the LFS). The LFS could induce a stabilized LTD in juvenile group but not adult group. This form of LTD could be drastically inhibited by GluN2B antagonist Ro25-6981 but not GluN2A antagonist NVP-AAM077. (\**P*<0.05, Juvenile or Juvenile+NVP group vs. other groups). **(D)** Example of filtered prelimbic LFP recordings of juvenile, adult and juvenile+anti animals during the PR-LTM test. Black lines, LFP filtered in the thta frequency range (8-12 Hz); gray line, LFP filtered in the gamma frequency range (30-100 Hz). Representative phase-amplitude diagrams (Top-right). The color scale represents the values of mean modulation index (MI). Juvenile rats showed a higher MI value than adult group while block of prelimbic proBDNF by its antibody dramatically reduced the MI value of juvenile group (Bottom-left). There was no statistical difference in the mean power of theta (Bottom-middle) or gamma (Bottom-right) oscillations. (\**P*<0.05, Juvenile or Juvenile+NVP group vs. other groups). The number of rats in each group used was indicated in each column/figure.

Figure 3C (bottom-left) showed the NMDA receptor-dependent *in vivo* LTD at hippocampal-prefrontal cortex synapses. Interestingly, the LFS did successfully induced LTD in juvenile but not adult rats. Herein, at the last 20 min of the recording, the average slope of juvenile group was significantly higher than adult and Juvenile+Anti groups (Figure 3C-(bottom-right); one-way ANOVA, *F*_(4, 22)_=3.13, *P*=0.035; post-hoc, Juvenile vs. Adult or Juvenile+Anti, both *P*<0.05). The LFS-induced LTD was completely suppressed by GluN2B antagonist Ro25 but not GluN2A antagonist NVP (post-hoc, Juvenile vs. Juvenile+Ro25, *P*<0.05; Juvenile vs. Juvenile+NVP, *P*>0.05), confirming a GluN2B-dependent synaptic plasticity. Additionally, input/output curves were established but the fEPSP slopes of juvenile rats did not differ significantly from adults (Supplementary Figure S2B). Meanwhile, LTD was also induced by a stronger LFS (900 pulses of 1.0 Hz) as previous methods (An *et al*, 2019; An *et al*, 2017, 2018b). The stronger LFS could induce robust LTD, which were comparable between juvenile and adult groups (Supplementary Figure S2C). Inactivation of proBDNF by its antibody could also effectively block the expression of the LTD in juvenile rats. These findings exclude the basal synaptic transmission effects on the LFS-induced LTD and further confirm a weaker LFS (300 pulses of 1.0 Hz) can induce LTD in juvenile but not adult rats. To assess whether and how ventral hippocampus is involved in updating fear memory and synaptic function, four-tone conditioned rats were infused with anti-porBDNF into ventral hippocampus immediately following memory retrieval, and PR-LTM and the LFS-induced LTD in the hippocampal-prefrontal cortex pathway was detected as above (Supplementary Figure S3A). We found the inhibition of proBDNF in the hippocampus did not alter retrieval-dependent memory destabilization (Supplementary Figure S3B). Anti-proBDNF antibody could dramatically depress 300-pulse LFS-induced LTD (Supplementary Figure S3C) but not 900-pulse LFS-induced LTD (Supplementary Figure S3D) in juvenile animals. These results clearly indicate that proBDNF in the ventral hippocampus did not play a critical role in retrieval-dependent fear memory destabilization or the expression of LFS-induced LTD.

Figure 4D (top) showed that the gamma oscillation of mPFC neurons tended to phase-lock to the peak of theta oscillation. Gamma amplitudes were coupled to theta oscillation during the PR-LTM test and that this modulation was stronger in juvenile group than in adult group (Figure D-(bottom-right); one-way ANOVA, *F*_(2, 14)_=7.53, *P*=0.006; post-hoc, Juvenile vs. Adult, *P*<0.05). Blockage of the increase proBDNF did reduce the phase-amplitude coupled theta and gamma oscillations compared to juvenile group (post-hoc, Juvenile vs. Juvenile+Anti, *P*<0.05). Furthermore, we found equalized theta (Figure 4D-(bottom-middle); one-way ANOVA, *F*_(2, 14)_=0.42, *P*=0.665) and gamma power (Figure 4D-(bottom-right); one-way ANOVA, *F*_(2, 14)_=1.13, *P*=0.351) during the PR-LTM test among groups, indicating that the difference in coupling strength reflected an active memory process but was not simply the result of improved phase identification due to the increment in power.

## 4. Discussion

Growing findings in younger rodents indicate that fear expression may vary across neurodevelopment associated with dynamic alterations in fear-related neural circuits. Memory modification using reconsolidation updating is being examined as one of the potential treatment approaches for attenuating memories associated with neurodevelopmental disorders (Beckers and Kindt, 2017; Liberman *et al*, 2006; Merikangas *et al*, 2010). However, the underlying mechanism is still unclear. The involvement of proBDNF in fear memory extinction and expression has been demonstrated. Our previous studies reported that proBDNF was highly expressed during the postnatal period and blocking proBDNF-mediated signaling induced spine development deficits in adult rodents. In this study, we further confirmed the expression levels of proBDNF and p75^NTR^ around the 4^th^ postnatal week were marked higher than the adulthood. Auditory fear memories conditioned with eight tones could induce stronger freeze behaviors than four-tone conditioning. This high expression level of prelimbic proBDNF enhanced the retrieval-induced destabilization of the four-tone conditioned weaker memory traces through activating p75^NTR^ and NMDA-GluN2B signaling. Exogenous proBDNF failed to promote the retrieval-dependent memory destabilization in adults but juveniles, indicating an aged-related specific effect of BDNF (Mattson *et al*, 2004; von Bohlen und Halbach, 2010). Additionally, infusions of proBDNF or its antibody into the juvenile prelimbic region did not change PR-STM or produce remote effects on PR-LTM. Dendritic morphology of prelimbic neurons found higher proportion of thin type spine than anti-proBDNF condition following memory retrieval. Meanwhile, high GluN2B but low GluN2A levels were observed in juvenile animals. The facilitation of neuronal function was verified as evidences by elevating the amplitude and decay time of the NMDA-mediated currents. The GluN2B- but not GluN2A-dependent LTD at hippocampal-prefrontal cortex pathway was enhanced following the memory retrieval. We also found prelimbic theta-gamma coupling of juvenile rats was significantly strengthened during the PR-LTM test. Inactivation of proBDNF could drastically inhibit retrieval-dependent memory destabilization, suppress synaptic depression and reduce the neural correlate to the similar level as adults, providing compelling evidence for the pivotal role of proBDNF in memory retrieval and the mechanism.

The expression proBDNF and p75^NTR^ in rodents is developmentally regulated with high levels in the early developmental period of life, correlating well with the timings of synapse formation and elimination (Bartkowska *et al*, 2010; Yang *et al*, 2009c). The negative retrograde effects of proBDNF on neuronal connectivity and cell survival have been strongly linked to the improvement in memory extinction and flexible behavior (Sun *et al*, 2020b; Sun *et al*, 2018a; Xu *et al*, 2011; Yang *et al*, 2009a). Remarkably, spine dynamics are associated with the spine tip where the spine is in contact with the presynaptic bouton (Bhatt *et al*, 2009; Kasai *et al*, 2021). In this study, we found blocking proBDNF expression during the juvenile period selectively elevated thin type spines but reduced mushroom ones following memory retrieval. It has been suggested that mushroom spines represent more stable memory spines, which are expected to maintain the fundamental feature of mature neural circuits (Bourne and Harris, 2007; Yang *et al*, 2009b). The thin spines formed early during development are highly motile and plastic and have been hypothesized to comprise a specialized subpopulation, distinguished by their contribute to synaptic reorganization and new learning (Bourne *et al*, 2007; Kasai *et al*, 2010). Our findings are consistent with previous findings that the rapid decline in extinction learning of young animals coincide with the dynamic reorganization of synaptic spine circuitry within the prelimbic region, but not in infralimbic or frontal association cortex (Pattwell *et al*, 2016). Crucially, the secreted proBDNF can promote spine maturation by sustained elevation of intracellular calcium influx (Mizoguchi *et al*, 2021; Orefice *et al*, 2016), which is important in supporting memory destabilization (Da Silva *et al*, 2013; Jarome *et al*, 2016).

Electroconvulsive shock or systemic drug administration given after memory reactivation induces the destabilization for the original memory (Milton *et al*, 2008; Nader *et al*, 2000; Wang *et al*, 2009). Reconsolidation involving protein synthesis is required for retrieved memories to persist. Infusion of the protein synthesis inhibitor anisomycin into the amygdala can return fear consolidated memories to a labile state (Nader *et al*, 2000). We found intra-prelimbic infusions of anisomycin and proBDNF shortly after training prevented consolidation of fear memories. These observations are generally agree with recent findings that blocking of the preforntal ERK/CREB-signaling pathway, which is able to be activated by both endogenous and exogenous BDNF, regulates the transition of memory phase from reconsolidation to extinction and this process functions as a switch that cancels reconsolidation of fear memory (Fukushima *et al*, 2021). However, this suppression was negligible in the stronger conditioned groups. It is possibly due to the fact that strong memories can be resistant to this approach. It has been believed that the strong training down-regulates GluN2B expression, thereby making the memory insensitive to post-reactivation anisomycin infusions but capable of being expressed normally (Ben Mamou *et al*, 2006; Wang *et al*, 2009). In fact, an increase in intensity of auditory fear conditioning training lead to an increase in the GluN2A/GluN2B ratio and that this increase is sufficient to prevent retrieval-dependent memory destabilization and therefore inhibits the induction of reconsolidation (Holehonnur *et al*, 2016). Further researches identified the distinct roles of NMDARs, indicating GluN2B-NMDARs being required for destabilization and GluN2A-NMDARs being required for restabilization (Milton *et al*, 2013). An increase in NMDA-GluN2B expression can shift the modification threshold to the left thereby leading a lower threshold for LTD induction (Paoletti *et al*, 2013). In consistent with this switch in NMDAR subunit composition during the developmental epoch (Liu *et al*, 2004), the higher GluN2B expression was found during the 3^rd^ postnatal week in this study. Previously, a strong NMDAR-mediated EPSCs was induced by the current threshold LFS protocol, leading calcium influx through NMDAR with GluN2B subunits was sufficient for reaching the threshold for LTD induction in infant (PD11-PD14) rats (Yasuda and Mukai, 2015) and in hippocampal slices infused with proBDNF (Woo *et al*, 2005). This functional recruitment during early development stage is profoundly influenced by the permissive role of proBDNF on shaping synaptic network, with the best example being the apparent increase in amplitude and decay time of EPSCs (Gibon *et al*, 2015; Mizoguchi *et al*, 2021; Yang *et al*, 2009a), making them more susceptible for the occurrence of plastic changes. In prevailing models, the level and kinetics of NMDAR activation determine postsynaptic AMPAR trafficking, which is the critical step lead to memory destabilization (Holehonnur *et al*, 2016; Hong *et al*, 2013). In addition, there is a consensus concerning the balance between neuronal excitation and inhibition regulating by proBDNF in basal excitability threshold at developing synapses (Deinhardt and Chao, 2014; Langlois *et al*, 2013; Riffault *et al*, 2018), implying that metaplasticity mechanisms may serve to alter the specific reactivation cues necessary to destabilize fear memory (Finnie *et al*, 2012; Zhang *et al*, 2018). This provides an interesting avenue for future studies.

BDNF-mediated synchronizes neural activity in the mPFC appears to be critical for extinction memory formation and modulation (Hallock *et al*, 2019; Hill *et al*, 2016; Rosas-Vidal *et al*, 2014). Emerging evidence indicates that gamma oscillatory activity is shown to be heterogeneous during extinction learning, with variations in the strength of gamma power (Courtin *et al*, 2014; Fenton *et al*, 2016; Mueller *et al*, 2014). Actually, the gamma amplitude is modulated by the phase of theta, and it is theorized that theta-gamma phase-amplitude coupling coordinates neural activity at the timescale required for memory processing (Benchenane *et al*, 2011; Nyhus and Curran, 2010). Previously, the strength gamma-theta coupling increased when animals evaluated choice-relevant information (Lundqvist *et al*, 2011; Roux and Uhlhaas, 2014), which was essential for the destabilization of fear memory at the moment of recall (Radiske *et al*, 2020). Impaired fear extinction in Bdnf-e4 mice is accompanied by decreased hippocampal-prefrontal theta phase synchrony during early extinction, as well as increased mPFC activation during extinction recall (Hill *et al*, 2016), implying that activity-dependent BDNF signaling is critical for regulating oscillatory activity, which may contribute to altered behavior. These behavioral associations may be facilitated in part by proBDNF-mediated oscillatory recruitment of functional neuronal ensembles, such as the firing of regular-spiking neurons during theta oscillations (Sun *et al*, 2018b). Furthermore, recruitment of neuronal ensembles to specific oscillations is thought to occur in part through subthreshold resonance properties of neurons (Stark *et al*, 2013). This hypothesis is in agreement with findings showing that endogenous proBDNF regulates neurons to amplify their response to the oscillatory phase coupling (An *et al*, 2018a; Sun *et al*, 2019). Additionally, the role of proBDNF in neural coherence is in keeping with the known roles in synaptic structure and subthreshold LFS induced LTD. Interestingly, enhancement in proBDNF-mediated spike-field coupling reflects the stabilization of striatum-dependent reversal learning, leading to facilitation of repetitive choices by bridging between previous and current choices (Sun *et al*, 2020b). Furthermore, functional connectivity within the neural circuitry mediating learned fear inhibition, which includes reciprocal projections between mPFC and amygdala, involves functionally communicate via synchronized neural oscillations (Chen *et al*, 2021; Likhtik and Paz, 2015). Intriguingly, 8 Hz stimulation of amygdala can induce amygdalo-cortical network states only following extinction learning (Ozawa *et al*, 2020), implying that post-extinction fear memory retrieval is supported by both local and interregional experience-dependent resonance.

In this study, we tested the hypothesis that proBDNF plays a vital role in synaptic function and memory destabilization and therefore, the activation of proBDNF signaling can be used to enhance erasure of reconsolidation-resistant fear memories. However, these actions from the temporal elevated proBDNF are probably limited to an early postnatal time window. Our finding showed that intra-prelimbic infusions of exogenous proBNDF failed to promote the destabilization of an existing fear memory, which was coincided with early findings in the different regulations of psychostimulant in BDNF signaling between juvenile and adolescent rats (Banerjee *et* al, 2009; Kozisek *et al*, 2008). Therefore, determining the role of proBDNF in mediating adult neural circuitry underpinning learned fear inhibition is another key issue to investigate. Local γ-aminobutyric acid (GABA) interneurons in PFC, which regulate learned fear and its extinction, also modulate theta and gamma oscillations locally (Chen *et al*, 2020; Piantadosi and Floresco, 2014). Importantly, the NMDA-dependent switch of proBDNF/p75^NTR^ signaling can orchestrate the excitatory/inhibitory developmental sequence leading to depolarizing and excitatory actions of GABA in adulthood (Langlois *et al*, 2013; Riffault *et al*, 2018). The mechanisms underlying this effect needed to be further determined.

Our findings strongly suggest that the mPFC is engaged in destabilization of consolidated memory during memory retrieval, and the high expression of proBDNF during the juvenile period facilitate this processing via activation of the p75^NTR^ and NMDA-GluN2B signaling. Furthermore, proBDNF-mediated synaptic and neural function is involved in regulation of retrieval-dependent memory. Our results also suggest that phase-amplitude coupling analyses from EEG signals recorded can be useful to verify the actual occurrence of retrieval process and predict the treatment’s efficacy. Therefore, elucidating the role of the proBDNF in fear memory modification improves our understanding of the molecular mechanisms that can regulate the induction of destabilization/reconsolidation of the stored memories, as suggested by the trace dominance theory (Eisenberg *et al*, 2003).

## Acknowledgements

This work was supported by Grants from the National Natural Science Foundation of China (32160196; 31700929) to LA.

## Ethics Statement

All animal experiments and procedures were reviewed and approved by the Experimental Animal Care Committee of Guizhou University of Traditional Chinese Medicine (SCXK-2013-0020).

## Data Availability Statements

The data that support the findings of this study will be available in Supplementary Files and other data are available from the corresponding author upon reasonable request.

## Author Contributions

Conceived and designed the experiments: WS, YY, XC, XLL and LA; performed the experiments: WS, YZM, XLL and XC; analyzed the data: WS, XLL and XC; wrote the manuscript: WS, YY, YZM and LA.

